# Modeling tumor transport and growth with poroelastic biopolymer networks

**DOI:** 10.1101/2025.09.23.678021

**Authors:** Zecheng Li, Yifei Ren, Sharvari Kemkar, Paul Mollenkopf, Jakub Kochanowski, Paul A. Janmey, Prashant K. Purohit, Ravi Radhakrishnan, Kyle H. Vining

**Affiliations:** Department of Bioengineering, School of Engineering and Applied Science, University of Pennsylvania, Pennsylvania, 19104, United States; Department of Mechanical Engineering and Applied Mechanics, School of Engineering and Applied Science, University of Pennsylvania, Pennsylvania, 19104, United States; Penn Institute for Computational Science, University of Pennsylvania, Pennsylvania, 19104, United States; Department of Chemical and Biomolecular Engineering, School of Engineering and Applied Science, University of Pennsylvania, Pennsylvania, 19104, United States; Department of Physiology, School of Medicine, University of Pennsylvania, Pennsylvania, 19104, United States; Department of Preventive and Restorative Sciences, School of Dental Medicine, University of Pennsylvania, Pennsylvania, 19104, United States; Department of Materials Science and Engineering, School of Engineering and Applied Science, University of Pennsylvania, Pennsylvania, 19104, United States; Center for Innovation and Precision Dentistry, University of Pennsylvania, Pennsylvania, 19104, United States

**Keywords:** Poroelasticity, Tumor growth, Alginate hydrogel, Growth factor transport, Tumor microenvironment

## Abstract

The mechanical properties of the extracellular matrix (ECM) regulate tumor growth and invasion in the tumor microenvironment. Models of biopolymer networks have been used to investigate the impact of elasticity and viscoelasticity of ECM on tumor behavior. Under tumor compression, these networks also show poroelastic behavior that is governed by the resistance to water flow through their pores. This work investigates the hypothesis that poroelastic properties regulate tumor growth. Here, alginate hydrogels with tunable ionic and hybrid ionic/covalent crosslinking are used as a model biopolymer system. Hydrogel stiffness, viscoelasticity, and stress relaxation behavior were characterized using stepwise axial compression. Among these properties, we find poroelastic fluid outflow dominates ECM stress relaxation, as the measured water flux was significantly affected under compression. Continuum mechanics-based modeling was developed to formulate and calculate the chemical potential gradients of water (solvent) in the hydrogels under compression. This framework was extended into an advection-diffusion framework to quantify growth factor (solute) distribution under varying strengths of stress and diffusion indexed by the relative strength of convective to diffusive transport, characterized by the Péclet number. An agent-based computational simulation showed that tumor growth was affected by Péclet number. Together, these results highlight the role of the poroelastic properties of ECM on water flux and transport in the tumor microenvironment.

## I. INTRODUCTION

The extracellular matrix (ECM) plays a critical role in regulating cell behavior, including proliferation, migration, and differentiation, through both biochemical and mechanical cues. Mechanical cues of ECM arise from the structure and properties of biopolymer networks and their interstitial fluid flow. Stiffness, viscoelasticity, and plasticity significantly impact cancer cell mechanobiology, as cancer cells sense and respond to variations in mechanical forces determined by these mechanical properties.^1–6^

Interstitial fluid flow within the network’s pores regulates cancer cell migration and transport of nutrients, including metabolites, proteins, and cytokines, which have downstream effects on cancer cell behaviors.^7–9^ Under deformation, both biopolymer network and interstitial fluid flow contribute to the ECM’s stress relaxation properties. Biopolymer networks relax stress via structural reorganization (viscoelasticity) and fluid efflux from the porous network (poroelasticity). Viscoelastic and poroelastic relaxations manifest at different timescales.^10^ Oscillatory shear rheology and stress relaxation with small deformation (≤ 10% strain) and short timescales (0.1-100s) are used to characterize viscoelasticity of biopolymer networks.^2,11,12^ However, the mechanical behavior of biopolymer networks under large uniaxial or biaxial deformation (>10%) and long timescales (1000 - 100,000s) remains understudied.

Tumor proliferation generates large deformation over long timescales and pushes radially outward on the surrounding tumor microenvironment (TME), leading to self-compression due to physical confinement.^5,13^ Compressive stress experienced by solid tumors impacts tumor growth dynamics, morphology, and therapy resistance, ultimately shaping disease progression. In responses to compressive stress, tumor cells intrinsically undergo glycolytic reprogramming, enter quiescence that can impair drug efficacy, and activate mechanosensitive YAP/TAZ, PI3K–AKT, and Wnt–*β* -catenin pathways, collectively enhancing survival and invasiveness.^14–17^ Solid stresses during compression can cause collapse of blood vessels and affect lymphatic drainage.^13,18^ Compression during tumor growth also generates a pressure gradient and outward convective flux.^19–21^ Solid-phase compression and interstitial fluid flow are intrinsically coupled by the poroelastic nature of the ECM. Under large-scale, slow deformation, poroelastic models show interstitial fluid flow that influences the availability and distribution of important solutes like oxygen, growth factors, nutrients, and cytokines mediating tumor-immune interaction.^15,22^ Thus, investigating the poroelastic properties of tumor ECM may provide new strategies to mitigate compression-driven resistance and improve treatment efficacy.^17^

We hypothesize that poroelastic properties govern tumor behavior at relevant scalings of tumor growth compared to the faster and shorter length-scales of viscoelastic properties.^23^ We developed a poroelastic experimental, theoretical, and computational framework for characterizing compression-induced fluid flux in advection-diffusion simulations to quantify how deformation-driven fluid flow modulates growth factor distribution. Alginate hydrogels provide a model system of TME matrix properties. Alginate provides a natural polysaccharide-based network that inherently demonstrates compressive strain-stiffening properties,^24,25^ which is a key property of native ECM.^26,27^ Alginate’s nanoporous structure closely mimics the compact environment of TME, which limits interstitial fluid flow.^21^ In this work, we used large (up to 40% strain) and slow compression (0.025%/s) on an alginate hydrogel to model ECM deformation and outward fluid flow within TME during tumor growth. Stepwise compression was used to characterize stiffness and poroviscoelasticity, revealing poroelasticity as the key changing property. A customized rheological setup and finite element analysis (FEA) were utilized to evaluate the volumetric flux and chemical potential gradient of fluid within the polymer network, providing a basis for fluid dynamics within the matrix for solute transport phenomena and tumor tissue agent-based modeling. Agent-based modeling is used to simulate the spatiotemporal dynamics of individual entities, or agents, interacting within a shared environment. In the context of solid tumor growth, each agent represents a cancer cell, which resides within a domain representing the tumor microenvironment (TME). Cancer cells proliferate based on user-defined growth rates.^28,29^ In this study, the agent-based model helps in the direct quantification of growth factor distribution-mediated differences in tumor cell growth. We used the computed growth factor concentration profiles under compression to simulate tumor growth dynamics using an agent-based modeling platform. Overall, our results offer key insights into the role of compressive stress-induced interstitial flow in ECM remodeling during tumor progression.

## II. RESULTS AND DISCUSSION

### A. Fluid flow dominates in biopolymer networks with large and slow deformation

Stress relaxation behavior reveals the mechanisms of ECM relaxation through its two components: the biopolymer network and the interstitial fluid. The biopolymer network relaxes stress via structural reorganization (viscoelasticity), while the interstitial fluid contributes through fluid transport and efflux from the porous network (poroelasticity)^23^. Stepwise compression, involving a series of 10% compressions, 2500s of stress relaxation, and time sweep after complete relaxation, was used to measure relaxation time constants, storage moduli, and tan(*δ*) (viscoelasticity). Relaxation time constants were obtained by fitting stress-relaxation curves to an exponential decay model based on the Generalized Maxwell Model, while storage moduli and tan(*δ*) were directly measured from time sweep rheology (Figure 1A). Ionic-crosslinked alginate (via calcium ions) and covalent-crosslinked alginate (via Nb-Tz click chemistry) were tested under stepwise compression to assess their relaxation time constants. After fast compression (strain rate of 0.25%/s) from 0 to 10% strain, the relaxation time constant of covalent alginate was approximately three times higher than that of ionic alginate, indicating faster relaxation in ionic alginate. Under larger strain compressions, covalent alginate exhibited distinct trends and significantly different relaxation time constants compared to ionic alginate (Figure 1B). These differences likely arise from the nature of crosslinks, as chemical hydrogels with rigid covalent bonds are less likely to break and rearrange and thus relax slower than physical hydrogels^23^. However, after slow compressions (strain rate of 0.025%/s), both covalent alginate and ionic alginate showed similar relaxation time constants and trends (Figure 1C), suggesting their stresses are released by a similar relaxation mechanism. Since the expected relaxation difference due to crosslink (biopolymer network reorganization) differences did not result in a prominent difference in relaxation time constant, biopolymer network reorganization plays a minimal role under slow deformations. This implies that fluid flow becomes dominant during large and slow compressions, as seen in processes of tumor growth and tissue development.

**FIG. 1.**
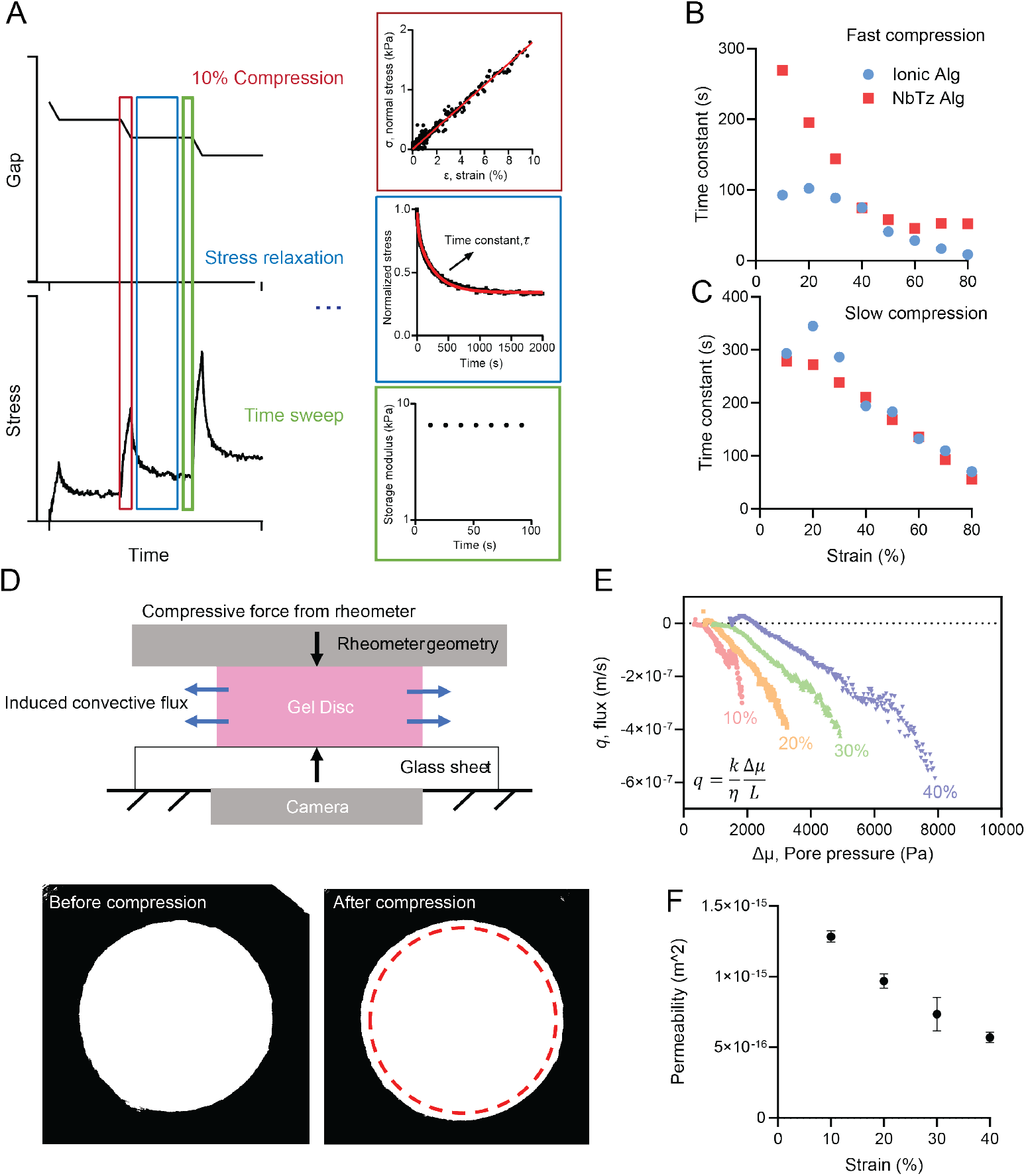
Fluid flow dominates stress relaxation behavior during large and slow deformations. A) Stepwise compression and stress relaxations with time sweep, which quantifies relaxation time constants, storage moduli, and tan(*δ*). B–C) Relaxation time constants at various strains for ionic and covalent crosslinked alginate (Alg) after fast, 0.25%/s strain rate (B) and slow, 0.025%/s strain rate (C) compressions. D) Compression on gel disc generates the outward convective flux, which is measured by a customized rheometer setup via recording volume change over time. E) Ionic alginate’s flux and pore pressure difference change during stress relaxation under 10–40% compressive strain. F) Permeability of ionic alginate as compressive strain increases.

To investigate the impact of compression on fluid flow and solute transport in ECM, both experimentally and computationally, we focused on a disc section of ECM modeled by an alginate gel. To experimentally measure fluid flow, a customized rheometer setup was used to obtain volume changes during compression (Figure 1D) of the alginate hydrogel, which is used to calculate volumetric flux of the fluid, *q*^30^. As compressive strain increases, fluid flux becomes increasingly negative, indicating fluid is leaving the matrix at a faster speed. The measured flux of 0–0.6 *µ*m/s (Figure 1E) matches in situ interstitial fluid speed in neoplastic tissue (0.5–0.6 *µ*m/s)^21^, suggesting our alginate hydrogel and testing system is a reasonable model for ECM in tumor microenvironment. Darcy’s law, 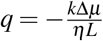, quantifies volumetric flux, *q*, of the fluid (with viscosity, *η*) moving through a matrix with permeability, *k*, under a pore pressure gradient, Δ*µ*, over distance *L*. Using Darcy’s law, ionic alginate’s permeability for each stress relaxation was determined by linearly fitting flux versus pore pressure difference (Figure 1E). The permeability of ionic alginate decreased with increasing compressive strain (Figure 1F), consistent with network densification. As the alginate disc is compressed and fluid is expelled, the polymer network becomes denser, making fluid permeation more difficult. Oscillatory shear rheology of the ionic gel was performed after the normal stress had fully relaxed (before each 10% compression step). Shear storage moduli and tan(*δ*) of the ionic alginate slightly increased, which was consistent with densification of the matrix (Supplementary Figure S1). These results together demonstrate the significant contribution of the poroelastic behavior of biopolymer networks under slow rates of compression.

### B. Water flux varies non-linearly as a function of compressive strain

To better understand compression-induced fluid flow dynamics, finite element analysis was conducted to compute the chemical potential (pore pressure) of the fluid within the network. The finite element model (FEM) was first constructed and validated using experimental stress-strain curves at three different strain rates (Supplementary Figure S2), which showed quantitative agreement between the model and experiments. The chemical potential, *µ*, is defined as *µ* = *p* − *π*, where *p* is the local hydrostatic pressure and *π* is the osmotic pressure. Since the hydrogel disc is confined between two impermeable plates, fluid can only exit through the lateral surface of the gel cylinder during compression. At the boundary (*r*_edge_), fluid escapes easily due to the absence of network confinement in the direction of flow. In contrast, fluid at the center (*r*_0_), remains trapped due to confinement by the polymer network in the radial direction (Figure 2A). As a result, the chemical potential varies with radial distance, forming a gradient, 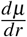, under compression. Volumetric flux at the boundary, *q*(*r*_edge_), can be calculated from Darcy’s law, 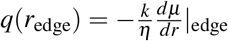, with the viscosity of fluid, *η*, and permeability, *k* (Figure 2B).

**FIG. 2.**
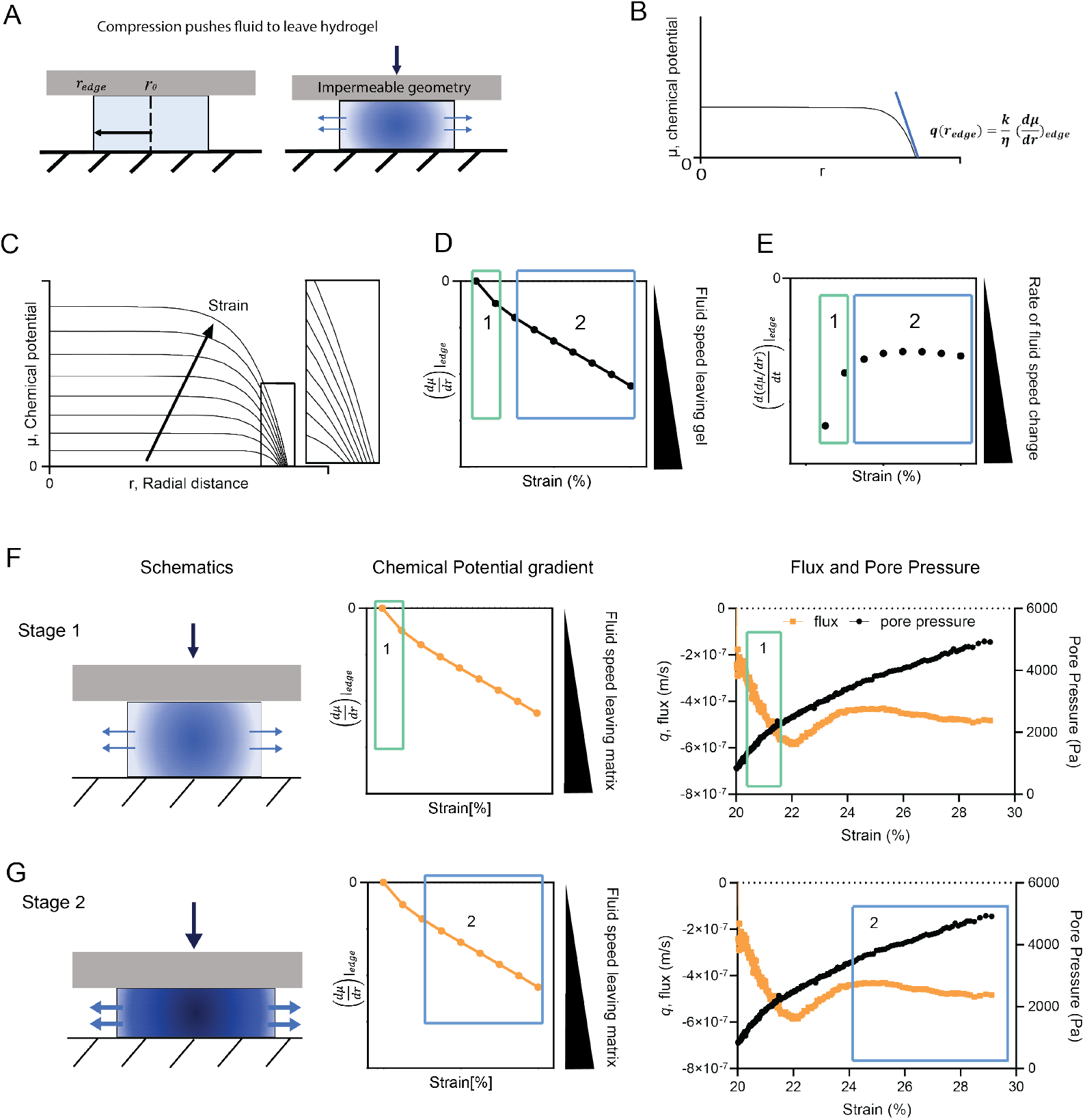
Nonlinear outward convective fluid flow under compression. A) Schematic of a high chemical potential fluid generated at the center of the hydrogel disc. B) Computational result: Chemical potential gradient under compression. C) Computational result: Chemical potential builds up as strain increases. D) Computational result: Change in chemical potential gradient at gel boundary, 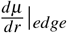, under increasing strain. E) Computational result: Rate of change in chemical potential gradient, 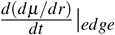, at gel boundary under increasing strains. F) Experimental result: Stage 1: Volumetric flux and chemical potential gradient get more negative faster, meaning fluid speed leaving the gel increases at a higher rate. G) Experimental result: Stage 2: Volumetric flux and chemical potential gradient decrease more slowly, meaning fluid speed leaving the gel increases at a lower rate.

As strain increases, fluid chemical potential builds up (Figure 2C). Due to the sudden generation of high chemical potential at the gel center from zero chemical potential at 0% strain, the chemical potential gradient at the boundary showed a fast decrease at the beginning of compression (stage 1) and followed by a slow and steady decrease at stage 2, showing a nonlinear water efflux under compression (Figure 2D-E). A more negative gradient indicates a greater chemical potential difference, leading to fluid leaving the hydrogel at a higher speed. The finite element simulation aligned with experimental findings, revealing two stages of fluid flux behavior during compression. In Figures 2F and 2G, fluid started with a fast decrease in flux at stage 1, then a more gentle and steady flux decrease at stage 2. Interestingly, two stages were also observed in stress-strain curves and correspond to two stages in flux change: when stage 1 transitioned to stage 2 for flux, fast-increasing pressure at stage 1 transitioned to the slow-increasing pressure at stage 2 as well (Figure 2F-G).

In stage 1, due to the sudden generation of high-pressure fluid at the center and its inability to immediately leave the gel, a large increase in chemical potential gradient is induced. As compressive strain increases with time, the high-pressure fluid in the center reaches the gel boundary and leaves without any network confinement, releasing the large difference in chemical potential gradient at stage 1 and entering stage 2. A transition region between stages 1 and 2 was observed (Figure 2F, Supplement S3). This phenomenon may result from the high-pressure fluid hitting the gel boundary and thus needing time to equilibrate and transition into stage 2. However, since stage 1 and transition stage only occur at the start of compressions and tumor growth is a long timescale process, we focused solely on the steady flux and chemical potential gradient change of stage 2 for the growth factor concentration profile and tumor growth simulation.

### C. Péclet number governs growth factor transport and distribution in the alginate gel geometry

Next, we modeled the effects of fluid flow on growth factor transport. Mitogenic growth factors are usually produced locally within the tumor microenvironment by tumor and stromal cells. Tumor cells acquire the ability to produce mitogenic growth factors for self-sufficiency, which is one of the two major hallmarks of cancer, along with the ability to sustain angiogenesis.^31–34^ Compression-induced stress influences both interstitial fluid flux and ECM permeability. Several studies have analyzed the impact of compression and permeability changes on solute convection and diffusion transport in TME, with the two processes showing different sensitivities to permeability changes based on the characteristics of the biopolymer networks and the solute properties.^9,35–37^ Since solute distribution depends on the relative strengths of convective and diffusive transport, we characterize the balance with the Péclet number in the advection-diffusion solute transport PDE modeling framework.

We simulated growth factor transport in the alginate gel subjected to 30 % compression over 5 h. Using the baseline run as a reference, we systematically varied the Péclet number to assess how changes in advective versus diffusive transport influence growth factor concentration fields. Figure 3 shows the ranges of Pe number corresponding to transport in our alginate gel domain of characteristic length *L* = 4 mm. The diffusion coefficient (*D*) of a typical growth factor, e.g. vascular endothelial growth factor or VEGF, (size 35-50 kDa) is set at 100 *µ*m^2^*/*s.^38^ The temporal chemical potential gradient profiles obtained (Supplement Fig S4) from simulating 30 % compression over a 5-hour period in the continuum finite element modeling framework, result in a maximum velocity (*v*) of ∼ 5.5 *µ*m/s (Supplement Fig S6). Given that permeability variations remain within a narrow range (Supplement Fig S5), we assume the diffusion coefficient to be spatially and temporally invariant for the purposes of this model. This gives us 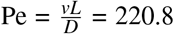 for the selected *D*−*v* combination. Assuming that permeability changes are minimal and the non-dimensionalized velocity profiles scale with compression under small strain rates in a fixed geometry, we utilize the same spatiotemporal profiles and scale them for simulating lower Pe cases at 0.2208, 2.208, 5, and 22.08. Note that if large compressive strains alter permeability significantly, the profile shape must be recomputed rather than merely scaled.

**FIG. 3.**
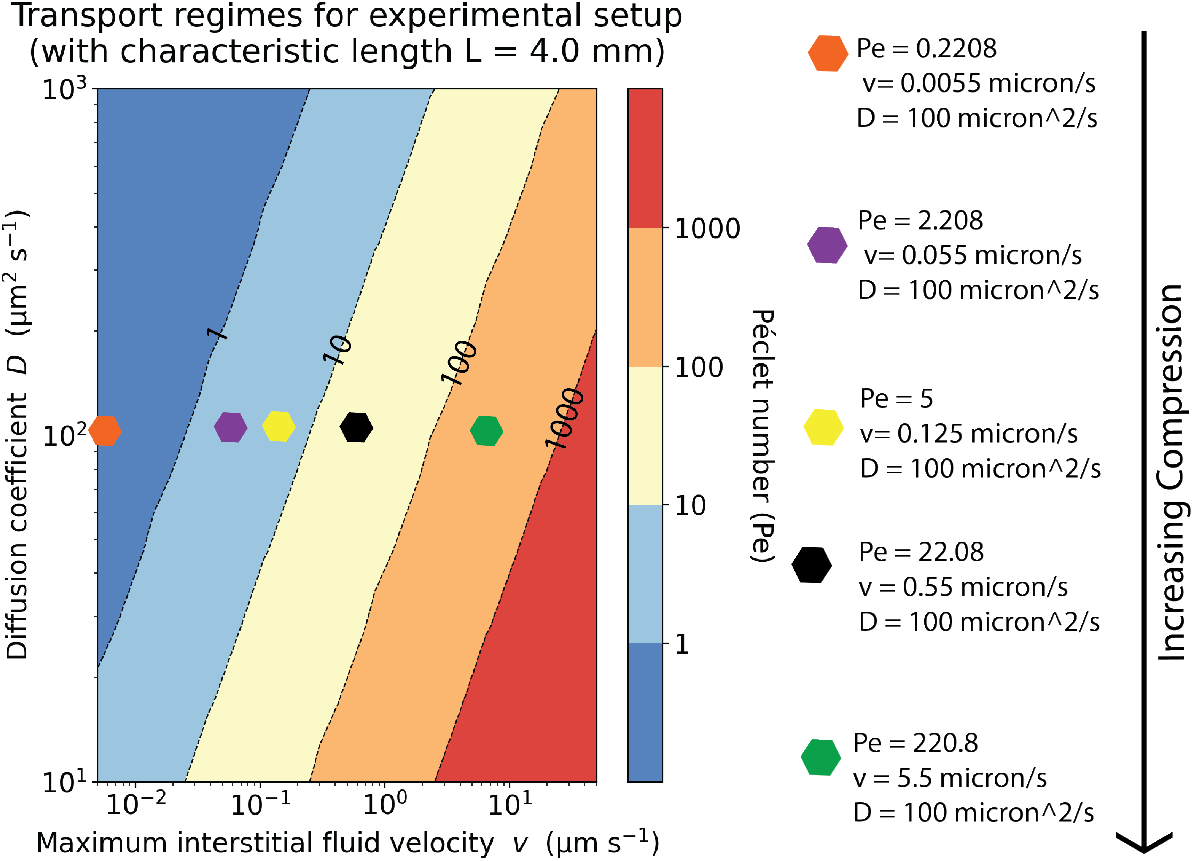
Physiological ranges of interstitial fluid velocity (*v*) and solute diffusivity (*D*) drawn from tumor and hydrogel studies were mapped to Pe values. We thus solved for three representative transport regimes: Pe = 0.2208, 2.208 (diffusion-biased), 5, 22.08 (mixed), and 220.8 (advection-dominated) to represent the spectrum of in vivo and in vitro TME transport behavior. The underlying *v* and *D* ranges and their primary literature sources are summarized in the methods section.

We modeled the concentration dynamics of a prototypical growth factor under two scenarios: 1) growth factor saturated domain - where the tumor microenvironment is initially saturated with growth factors; initial concentration (non-dimensionalized) of the growth factor set to 1 and the boundary condition across all domain edges is set to 0 (Dirichlet boundary condition); 2) growth factor depleted domain - where the tumor microenvironment initially lacks growth factors, with growth factors supplied from the surrounding regions (e.g. blood vessels), diffusing into the tumor microenvironment; initial concentration of growth factor set to 0 and Dirichlet boundary conditions set to 1. The PDE setup, non-dimensionalization, and solution methodology have been outlined in the methods section. In the results section, we only show results from the growth factor saturated domain scenario. The growth factor depleted domain simulations and results have been discussed in the supplementary information (Supplement S8, S9).

In the growth factor saturated domain simulation, two transport processes operate simultaneously in radially outward directions: (i) passive diffusion driven by the emerging concentration gradient carries the growth factor radially outwards, and (ii) a radially outward interstitial fluid flux (generated by compression) carries the growth factor with it (Figure 4 A). At Pe = 0.2208, diffusion dominates, and the core concentration reduces to 15% of the maximum attainable concentration *C*_0_ in 6 h, with gradients penetrating deep into the domain. In the mixed-transport regimes (Pe = 2.208 and 5.0), advective flux begins to counterbalance diffusion, such that the central concen-tration remains near 0.4 *C*_0_ and 0.7 *C*_0_, respectively, at 360 min. At Pe = 22.08, the profiles are almost flat in the core at *≈* 0.95 *C*_0_, with depletion confined to a narrow boundary layer. Finally, under advection-dominated conditions (Pe = 220.8), the concentration remains essentially uniform (*C*^*∗*^ *≈* 1) throughout the entire domain over the full 360 min. Together, our numerical solutions reveal that the Pe number governs both solute retention/availability and spatial distribution in the domain. As the Pe number increases from 0.2208 to 220.8, the concentration profiles transition from diffusion-dominated decay to advection-dominated retention. Higher Pe simulations resulted in flatter concentration fields throughout the tumor microenvironment, thereby democratizing the regional availability of growth factors throughout the domain (Figure 4 B-C).

**FIG. 4.**
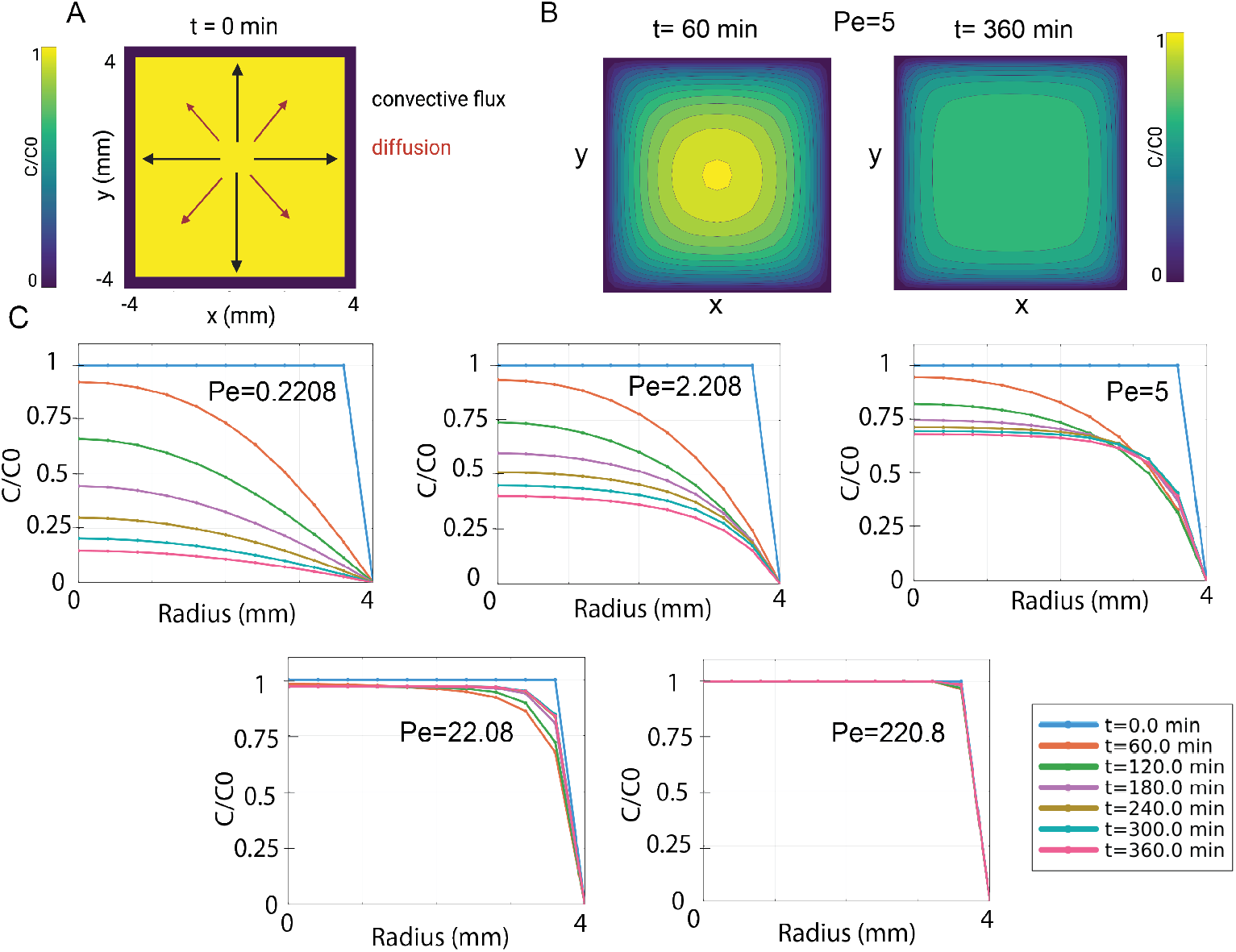
Influence of Péclet number on growth-factor distribution. A) At *t* = 0 min, the concentration field is uniform 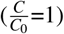 across the 8mm*×*8mm domain and boundary conditions are set to 0. Black arrows indicate the imposed outward convective flux, while red arrows show the direction of diffusion. B.) Two-dimensional concentration contours at *t* = 0 min (left) and *t* = 360 min(right) illustrate how the initially uniform field evolves under a moderate flow regime (here, Pe = 5.0). By 360 min, a nearly square-shaped plateau of high concentration persists in the interior, bounded by steep boundary layers. C.) Radial profiles of 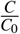 at six time points (0, 60, 120, 180, 240, 360 min) for five Péclet numbers (0.2208, 2.208, 5.0, 22.08, 220.8). Low Pe leads to significant interior depletion by diffusion, whereas raising Pe progressively preserves interior growth-factor levels and confines depletion to narrow boundary regions.

### D. Growth factor concentration gradients modulate tumor proliferation dynamics

Next, we quantified the difference in tumor growth propensities due to the spatial heterogeneity in access to growth factor levels by simulating tumor growth in an ABM. In our model, the primary source of spatial heterogeneity is manifested through a gradient in growth factor (solute) concentration due to the chemical potential gradient of water (solvent) established by poroelastic effects. Other sources of heterogeneity such as non-uniformity in the matrix or spatial effects in the stroma are not considered in our base model and can be incorporated in future extensions of our ABM. The Péclet number strongly influences the local availability of growth factors to tumor cells. In low-Pe regimes, diffusion dominates, leading to systemic depletion of growth factors and distinct spatial heterogeneity, whereas in high-Pe regimes, advective transport preserves elevated, more uniform growth factor concentrations throughout the domain. Growth factor concentration levels were used as input to the tumor ABM in PhysiCell, systemically varying 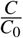 from 0 to 1. This framework was used to generate a family of tumor growth curves as a function of growth factor levels. Figure 5 summarizes the observed tumor growth dynamics over 7 days in a small 0.4 mm *×* 0.4 mm tumor microenvironment subsection under different growth factor availability. When the environment is replete (1) with growth factors or moderately supplied (0.64, 0.32), the cell population grows exponentially. Intermediate growth factor availability (0.16) yields slower growth, which is approximately linear. Below a threshold, net proliferation ceases: the 0.08 case remains nearly steady, while lower growth factor levels show a progressive decline in growth.

**FIG. 5.**
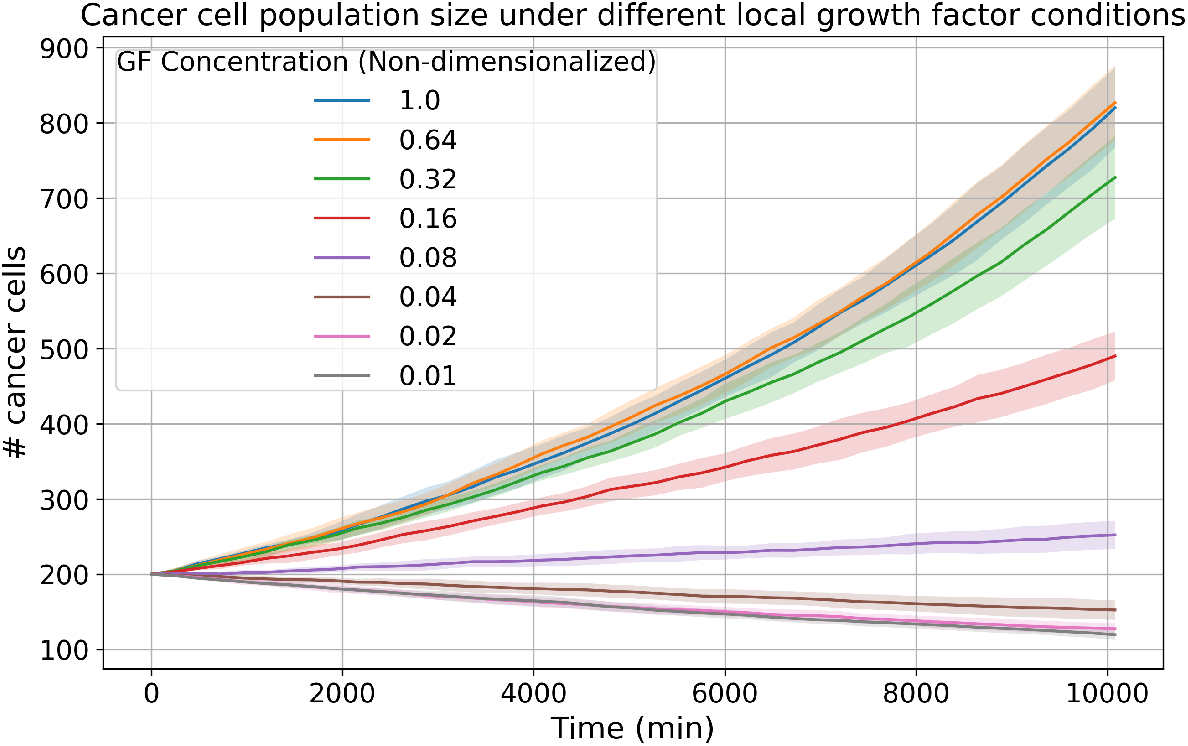
Temporal profiles of tumor load (cancer cell population count) - Family of curves generated from ABM simulations under varying growth factor levels. The listed growth factor levels are used as initial and boundary conditions for the PhysiCell ABM. Shaded envelopes indicate inter-replicate variability, which widens with higher growth-factor availability due to amplified stochastic proliferation. Overall, tumor load scales monotonically with growth factor availability, exhibiting net growth above 0.08 and net regression below it.

To compare tumor growth under different Péclet number conditions and spatial regions, we also conducted the ABM simulations using growth factor profiles in two distinct regions in our computational grid (Region A at *r* = 0 mm and Region B at *r* = 3.6 mm), under different Péclet number conditions. The advection-diffusion PDE is solved on an 8 mm *×* 8 mm domain, while an agent-based model (ABM) probes two 0.4 × 0.4 mm grids (Regions A and B). (Figure 6A). The growth factor availability at Regions A and B under different Pe values has been summarized in a heatmap (Figure 6B). The observed tumor load (total number of cancer cells) after a 7-day tumor growth ABM simulation under the corresponding growth factor availabilities has been summarized in a bar plot(Figure 6C). The cell counts remain low when Pe = 0.2208 and increase two to three-fold at higher Pe values. Under diffusion-dominated conditions (Pe = 0.2208, 2.208), mean tumor load at *t*=7 days for Region A was significantly higher than Region B. (*p <* 0.0001) Thus, Region A tumors grew significantly more than Region B tumors. However, at higher Péclet numbers, no significant difference in mean tumor load was observed between Regions A and B.

**FIG. 6.**
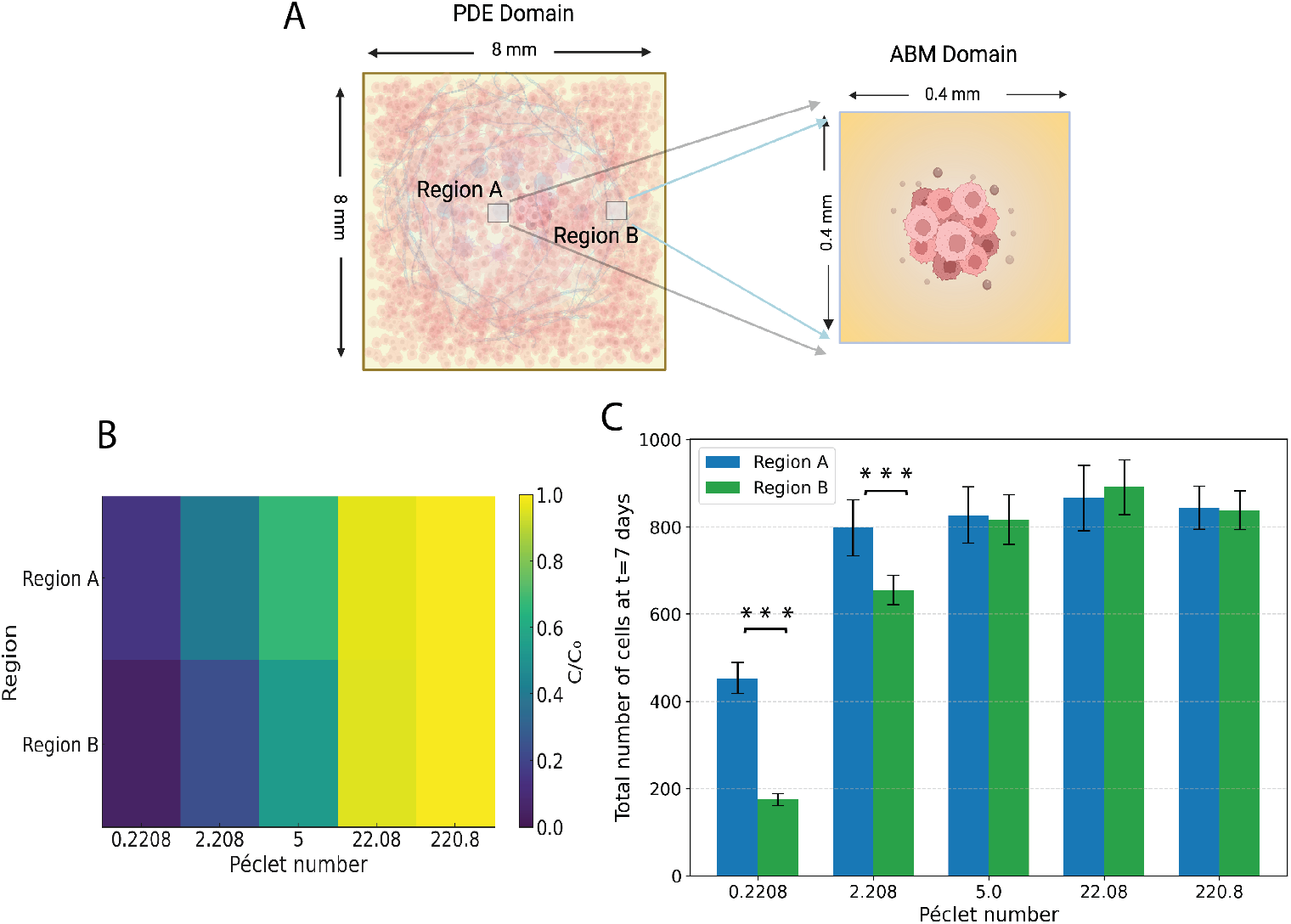
Coupling growth factor transport and tumor growth results across scales A) Setup of the compu-tational study - PDE domain on the 8 mm *×* 8 mm domain and tumor growth ABM domain on 0.4 mm *×* 0.4 mm regions of the domain. B) Heatmap - summarizing the Region A and region B growth factor (GF) concentrations (at *t*=6 h). Spatial heterogeneity in growth factor availability between Regions A and B reduces as Pe increases. Under high Pe conditions, growth factor levels are both elevated and more uniform across the domain. C) Tumor load (total number of cancer cells) at *t* = 7 days / 10080 min at Regions A and B, under different Pe. Tumor load bars show the *µ ± σ* from ten independent ABM stochastic runs per condition. Welch’s two-sample t-test was applied to each Region A-B pair; a single asterisk indicates a significant difference with *p* < 0.01 and three asterisks indicate *p* < 0.0001.

## III. DISCUSSION

By measuring fluxes under stepwise compression, we demonstrated that mechanical relaxation from fluid flow dominates viscoelastic reorganization in biopolymer networks undergoing large, slow deformation, such as those seen in solid tumor growth. (Figure 1B-C) This is further supported by the continuous decrease in permeability (Figure 1D-E) during compression and minimal changes in viscoelasticity and shear moduli (Supplement S1) below 40% deformation. Previous studies showed interstitial fluid flow regulates cell migration^8^ and solute transport^9^, while viscoelasticity regulates cell invasion^2^ and proliferation via short lengthscale mechanotransduction^1,3^. Our work suggests that fluid flow (poroelasticity) and solute transport dominate cell behavior in large and slow deformation of biopolymer networks.

We identified a non-linear behavior of fluid flux during compression with a customized rheometer and finite element modeling. Both experiments and simulation show that fluid experiences a fast-changing stage 1, and then a slower and steadily changing stage 2 (Figure 2D-G), which is interestingly reflected in the corresponding two stages observed in stress-strain curves (Figure 2F-G). In stage 1, due to the sudden generation of high-pressure fluid at the center and its inability to immediately leave the gel, a fast and large change in chemical potential gradient is induced. As strain increases, the high-pressure fluid in the center reaches the gel boundary and leaves without any network confinement, releasing the large difference in chemical potential and entering stage

2. Interestingly, a transition region (Figure 2F, Supplement S3) may result from the high-pressure fluid impacting the gel boundary, which requires time to equilibrate to stage 2. However, since stage 1 and the transition stage only occur at the start of compressions and tumor growth is a long timescale process, we focused solely on the steady flux and chemical potential gradient change at stage 2 for the following concentration profile and tumor growth simulation. A previous study identified the non-linear stress-strain relationship to fluid flow based on experiments and a theoretical model^39^. Our work expanded on this finding to show a non-linear flux behavior during compression by experiments and FEM simulation.

Previous computational models have shown that growth-induced solid stress and interstitial fluid pressure can influence tumor growth by suppressing proliferation, collapsing vessels, and inducing hypoxia.^16,40,41^ Liedekerke *et al*. developed an agent-based model for tumor cell growth by calibrating model parameters to mouse colon carcinoma compression experiments, using a strain-dependent Hill function for cell cycle progression.^42^ However, most studies assume or experimentally deduce a functional dependence of cellular proliferation/ tumor growth on compressive mechanical forces, without explicitly accounting for compression-induced mitogen transport in the tumor microenvironment. In this study, we coupled poroelastic mechanics to an advection-diffusion framework that explicitly computes compression-induced interstitial fluid flow and its effect on growth factor (solute) gradients. We studied this system under different Péclet number conditions and observed the spatial variations in the solute distribution. In the growth factor saturated environment, higher Péclet number conditions help retain more growth factors and promote more spatially uniform distribution of growth factors as compared to low Péclet number conditions (Figure 4 C). We employed an agent-based model to simulate and compare tumor growth under different Péclet number conditions. Our results show that Péclet number significantly impacts tumor progression. In a growth factor-saturated tumor microenvironment, high Péclet number (higher compression) promotes tumor growth, while low Péclet number results in slower progression or even regression of tumors (Figure 6 C). Under high Péclet number conditions, growth factors are more evenly distributed, resulting in similar local tumor growth rates across different regions of the tumor microenvironment (as shown by Figure 6 C high Pe cases, where no significant difference in mean tumor load was observed between regions A and B). In contrast, under low Péclet number conditions, steep solute concentration gradients emerge, resulting in heterogeneous growth factor availability and regionally variable tumor growth (as shown by Figure 6 C low Pe cases, where mean tumor load was found to be significantly different between regions A and B). Thus, Péclet number regulates spatial differences in solute availability, which directly contributes to region-specific differences in local tumor growth within the tumor microenvironment.

## IV. CONCLUSIONS

We have demonstrated that poroelastic fluid flow dominates biopolymer network behavior under large, slow compression. Using alginate hydrogels as an ECM model, we showed experimentally and computationally that interstitial fluid transport—rather than network reorganization—drives stress relaxation at long timescales relevant to tumor progression. We observed a nonlinear two-stage fluid flux behavior under compression governed by evolving chemical potential gradients and permeability that mirrors corresponding changes in stress-strain behavior. The advection-diffusion simulations of growth factor transport and tumor growth agent-based models were parameterized by the Péclet number. We observe that tumor growth is strongly influenced by the Péclet number. In growth factor-saturated conditions, high Péclet number promotes proliferation, while low Péclet number leads to slower growth or regression. Local region-specific agent-based simulations showed that low Péclet number leads to spatial heterogeneity in tumor proliferation, with higher tumor load accumulating in regions with higher growth factor availability. In contrast, high Péclet number conditions homogenize growth factor availability, which promotes spatially homogeneous tumor proliferation in the tumor microenvironment. We expect these compression-induced changes in transport may also affect immune responses through the lymphatic drainage of tumor antigens and cells, as well as immune infiltration and activation through the availability of chemokines and cytokines. Together, these results suggest that compressive stress exerted on biopolymer networks affects tumor growth through fluid-driven growth factor transport.

## V. MATERIALS AND METHODS

### A. Overall experimental and computational workflow

Flux and permeability were measured by compressing an alginate gel in a custom setup to model compression of the surrounding ECM during expansive tumor growth (Figure 7). Continuum-mechanics modeling and finite-element analysis were then used to map different compression levels to outward water flux, based on the temporal profiles of the chemical potential gradient of water. To explore how this compression-induced growth factor efflux affects tumor growth over longer timescales, we considered tissue under 30 % compression. We modeled growth factor advection–diffusion to obtain the resulting spatial distribution and simulated agent-based models of tumor growth under growth factor concentrations. In order to capture the effect of compression on growth factor distribution and tumor growth, we ran a sensitivity analysis on Péclet number by varying the effective maximum velocities associated with compression-driven water flux.

**FIG. 7.**
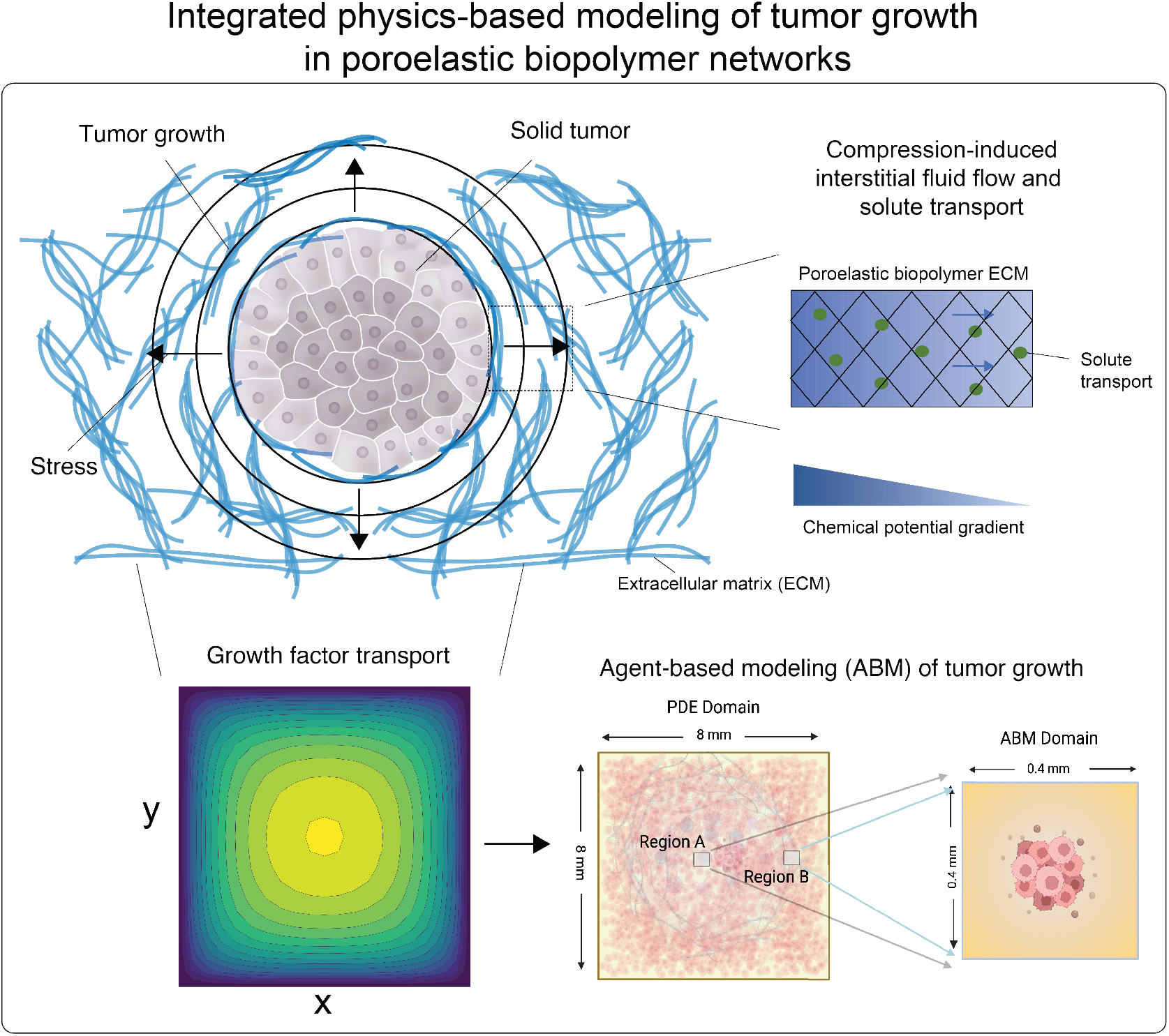
Schematic of the multiscale theoretical and computational framework of tumor growth in poroelastic biopolymer networks

Below, we describe each method in detail:

### B. Alginate Functionalization

Ultra-pure sodium alginate with very low viscosity (ProNova UP VLVG, NovaMatrix) was used, which we will be referred as VLVG in the remainder of the text. VLVG has an approximate molecular weight of 32 kDa and a guluronate-to-mannuronic monomer ratio of ≥ 1.5. VLVG is covalently modified by conjugating either 5-norbornene-2-methylamine (Nb, TCI America) or (4-(6-methyl-1,2,4,5-tetrazin-3-yl)phenyl)methanamine hydrochloride (Tz, Karebay Biochem), generating Nb-VLVG and Tz-VLVG, respectively.^11,25^ A theoretical degree of substitution (DS) of 25% was targeted. Initially, VLVG was dissolved at a 1% w/v concentration in a 2-(N-morpholino)ethanesulfonic acid (MES) buffer with 0.3 M NaCl at pH 6.5, adjusted using NaOH. Next, N-(3-dimethylaminopropyl)-N’-ethylcarbodiimide hydrochloride (EDC) and N-hydroxysuccinimide (NHS) were added at a 2.5-fold molar excess relative to alginate carboxylic groups and continuously stirred at 4^*°*^C. Nb or Tz was then added dropwise at a concentration of 4.7 mM per gram of VLVG. EDC, NHS, Nb, and Tz were divided into four aliquots, which were introduced at four intervals, each spanning two hours. The reaction continued for 18 hours at 4 °C with stirring. After completion, the solution underwent four rounds of ultracentrifugation at 14,000 rpm for 15 minutes each, followed by filtration through a 0.22*µ*m membrane. The product was then dialyzed using a 3.5 kDa molecular weight cutoff membrane (Spectra Por 3, Spectrum Labs) against a stepwise NaCl gradient of 150, 100, 50, and 0 mM in Milli-Q water over four days, with the buffer replaced twice per concentration each day. The final product was lyophilized and stored at -80^*°*^C.

### C. Fabrication of alginate hydrogels

VLVG alginate was prepared as previously described.^11,25^ A buffered salt solution was prepared by adding 20 mM 50x N-2-hydroxyethylpiperazine-N-2-ethane sulfonic acid (HEPES, Gibco) into 1x Hank’s Balanced Salt Solution (HBSS, Gibco) and adjusting the pH to 7.5 with 10 M NaOH. Stock solutions of unmodified alginate (VLVG) and amine-functionalized alginates (Nb-VLVG and Tz-VLVG) were prepared at a concentration of 5 wt% in HBSS/HEPES buffer. Precipitated calcium carbonate (CaCO_3_; Specialty Minerals) was dispersed in Milli-Q water at a 10 wt% concentration and ultrasonicated at 50% amplitude for 15 seconds to form a slurry. Freshly prepared D-(+)-gluconic acid *δ* -lactone (GDL; Sigma-Aldrich) was dissolved at 0.4 g/mL in HBSS/HEPES buffer immediately before use. For purely ionically crosslinked hydrogels, HBSS-HEPES buffer, CaCO_3_ with a concentration of 30 mM, VLVG with a concentration of 1.5wt%, and GDL solution (CaCO_3_:GDL molar ratio of 0.3) were sequentially added and fully stirred on ice. For click-modified alginates (Nb-VLVG and Tz-VLVG), the HBSS/HEPES buffer, CaCO_3_, Nb-VLVG stock solutions, and GDL solution with the same concentrations as unmodified VLVG alginate, were added sequentially with thorough mixing before introducing the Tz-VLVG stock solution. The click-modified alginate was made with an Nb-VLVG : Tz-VLVG ratio of 1:1, where covalent crosslinks were established via Nb-Tz click chemistry and ionic crosslinking was achieved with Ca^2+^.

### D. Modulus and relaxation characterization with stepwise compression

Hydrogel solution prepared as described above was loaded in a customized acrylic mold and polymerized for 5 hours at room temperature to form a hydrogel disc (*D × H* = 8 mm *×* 2 mm). The hydrogel disc was preconditioned for tests by soaking in Milli-Q water for 24 hours. Gel disc was transferred to a stress-controlled rheometer (DHR-30, TA instrument) and confined between two impermeable plates (top: 20 mm steel geometry; bottom: immersion cup) by lowering the top plate in contact with the hydrogel surface. Then, Milli-Q water was added to submerge the hydrogel disc and conditioned until reaching stable axial force. Stepwise compression was composed of 10% compressions followed by 2500 s of stress relaxations, ending after 8 steps and reaching 80% strain total compression. Elastic modulus was determined by performing a linear fit on the stress-strain curve for each 10% compression. Relaxation time constant for stress relaxation was characterized by fitting the stress-relaxation curve to an exponential decay function based on the Generalized Maxwell Model.

### E. Flux and permeability measurement with customized compression setup

The preconditioned hydrogel disc was transferred to a stress-controlled rheometer (DHR-20, TA Instruments) and positioned between two impermeable plates (top: 20 mm steel geometry; bottom: glass sheet placed above a camera). Once the geometry fully contacted the top surface of the gel, a Moticam S20 high-resolution camera was used to capture the stepwise compression described above, ending with 40% strain compression. The recorded video was pro-cessed through ImageJ to quantify the cross-sectional area of hydrogel disc over time. Assuming *V* (*t*) = *πr*(*t*)^2^*h*(*t*), where *πr*(*t*)^2^ is the cross-sectional area of the gel disc and *h*(*t*) is the gel thickness at time *t*, hydrogel volume, *V* (*t*), was calculated with *πr*^2^ obtained from ImageJ analysis and *h*(*t*) from the rheometer. The volumetric flux, *q*(*t*), is a measure of flow rate per unit area and is mathematically expressed as 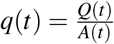 with unit m/s. Volumetric flow rate, 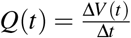, denotes the volumetric flow rate in unit m^3^/s. The fluid flow area, *A*(*t*) = 2*πr*(*t*) *· h*(*t*), denotes the effective area available for fluid outflow, where we assume gel with cylindrical geometry based on experimental observations. Permeability is an intrinsic parameter and characterizes porous material’s ability to allow fluid to flow through, which can be characterized by Darcy constant, *k*, with unit m^2^. *k* is calculated from Darcy’s law 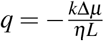 with viscosity, *η*, of fluid traveling over distance *L*, where a chemical potential gradient Δ*µ* will cause fluid flow. A Python script was used to synchronize and process both the stress data from the rheometer and the video-flux data, generating a curve of flux vs. normal stress (Figure 1E). Considering the equation for the Darcy permeability, a slope, 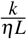,is obtained to calculate permeability *k* by fitting a linear function to the curve. By taking *L ≈ r*(*t*), we can approximate the chemical potential gradient 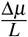 as 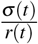, where *σ* is the average normal Cauchy stress on the top plate. We did so by following^4344^ who give a simple expression for fluid flow through compressed open cell foams in terms of the measured compressive stress and dimensions of the foam. We fitted the last linear part of the curve at the low flux region for each compression-relaxation step, because at the later phase of relaxation step the chemical potential gradient in the radial direction can be approximated as linear in agreement with our assumption above^30^.

### F. Continuum modeling and finite element analysis

#### 1. Kinematics

We focus here on the poro-elastic modeling of the alginate gel compression experiments. The gel is modeled as a continuum. Each material point in this continuum consists of a mixture of solid and fluid molecules and is considered homogeneous and isotropic in the initial undeformed configuration. Upon (large) deformation, the solid network can become anisotropic with different tangent moduli in different directions, governed by the local deformation field. Letting **x**(**X**, *t*) and **X** denote the current and initial positions of a specific material point, respectively, we can define the deformation gradient tensor and its determinant as:

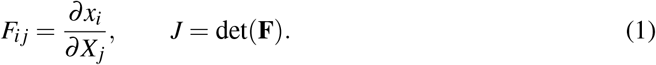

The initial (reference) configuration is taken to be the swollen state that the alginate network forms with water at equilibrium. We assume that the alginate material and the interstitial liquid are both incompressible and that volume change of the gel is caused purely by the movement of liquid. Let *C*(**X**, *t*) denote the volume fraction of liquid in the reference state, then:

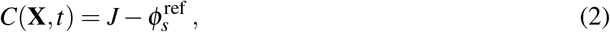

where 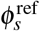 denotes the volume fraction of the polymer in the reference configuration. Now, if we consider a volume *V* with a normal **N**(**X**) and surface **S**(**X**) in the reference configuration, then mass balance in the absence of any liquid source can be written as:

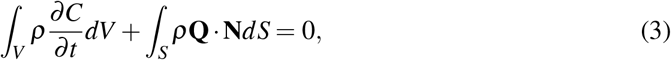

where **Q** denotes the liquid flux per unit reference area and *ρ* denotes the density of the liquid. Utilizing the divergence theorem, we can rewrite this equation in localized form as:

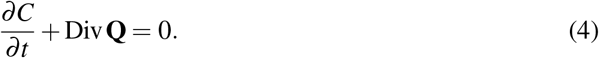

If we consider the loading to be quasi-static, which means that the load is applied slowly, we can write our momentum balance equation expressed in the reference configuration as:

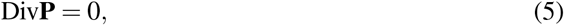

where **P** is the first Piola–Kirchhoff stress tensor. This type of chemo-elastic formulation has been adopted in previous studies^45,46^ and it is equivalent to the theory of porous media (TPM), including under large deformations. TPM explicitly accounts for the relative motion between solid and fluid by formulating separate equilibrium and mass balance equations for the solid and fluid constituents. It also incorporates body forces exerted on both the solid and fluid phases by the relative motion. It has been shown by Stracuzzi et al.^47^ that for incompressible solid and fluid constituents, the more detailed TPM framework is mathematically equivalent to the chemo-elastic formulation employed here.

#### 2. Constitutive laws

Experimental results indicate that the viscous effects are negligible and less important for long time scales and large compressive strains. Consequently, dissipation is caused only by the flow of liquid through pores and we consider the network to be elastic.Based on Flory-Rehner theory^48^, the total Helmholtz free energy per reference volume of the polymer network and liquid mixture can be written as a summation of the free energy density due to the deformation of the network Ψ_net_ (**F**), and the free energy density of interaction of liquid and solid Ψ_int_ (*C*):

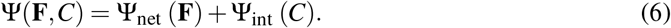

Utilizing Legendre transformation, we define a new free energy density as:

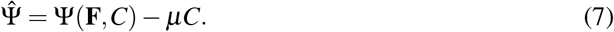

Here 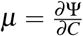 is the chemical potential of the interstitial liquid^46^. Based on standard continuum mechanical derivations that use the dissipation inequality, it can be shown that:

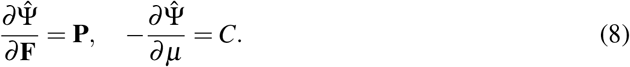

The free energy Ψ_net_ (**F**) and Ψ_int_ (*C*) used in this work is based on^47,49,50^:

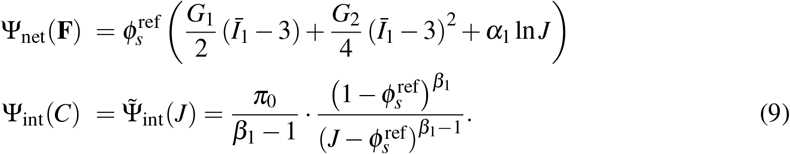

Notice that *C* in eqn. (9) is replaced by *J* according to eqn. (2). Here *G*_1_ and *G*_2_ are two constants. *G*_1_ is the small strain shear modulus of the dry polymer, while *G*_2_ is related to the shear modulus at large strain. A two-term polynomial hyper-elastic model is used to characterize the strain-stiffening behaviors of the polymer under compression. A ln *J* term is included in the expression for the free energy density following standard texts in nonlinear elasticity (e.g.,Ogden^51^, Holzapfel^52^) and biophysics (e.g., Boal^53^). 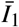 is the modified invariant of the left Cauchy-Green tensor **B**. The isochoric part of *I*_1_ is chosen to account for the incompressibility of both liquid and solid phases. *π*_0_ is the initial osmotic pressure, and the value of *α*_1_ is chosen such that the reference state is stress free. *β*_1_ is a power law coefficient, which is slightly larger than 1 in the current formulation. Utilizing eqn. (9), the Cauchy stress can be calculated as:

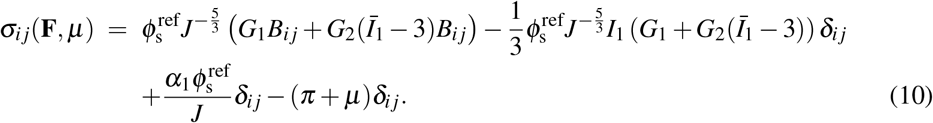

In order to set the value of *α*_1_, we require *σ* (**I**, 0) = 0 at *t* = 0, so that:

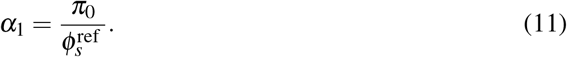

*G*_1_ and *G*_2_ are fitted to experimental stress-strain data from compression experiments as shown in the Supplement.

#### 3. Liquid flux

In this work, Darcy’s law is used for the constitutive behavior of the liquid flux. Further, we assume the material is isotropic so that this relation can be written as:

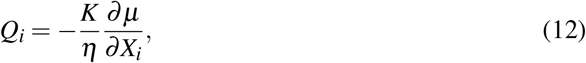

where *η* is the viscosity of the liquid and *K* is the permeability. *K* is a decreasing function of the current solid phase volume fraction *φ*_*s*_ which evolves with local deformation. We use the Carman-Kozeny expression for the permeability^54^:

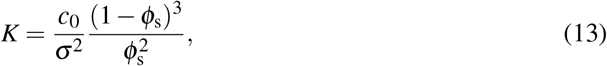

where *c*_0_ is a constant less than 1 and *σ* is the specific surface area. Here we assume that the alginate chains form tubes with circular cross section, which results in *c*_0_ = 0.5 and 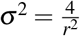. Note that the current solid phase volume fraction is related to the reference solid volume fraction by 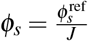, which depends on the local deformation.

To solve the full boundary value problem we use (a) impermeable boundary conditions on the top and bottom plates and permeable boundary conditions on the lateral surfaces, (b) displacement boundary condition on the top surface, fixed boundary condition on the bottom surface, and traction free boundary conditions on the lateral surface. The model is implemented in commercial finite element software ***ABAQUS*** Standard in the SOILS module with constitutive laws input through a user subroutine. C3D8P brick (Taylor-Hood) elements are used for the discretization.

### G. Growth factor transport

#### 1. Advection-diffusion PDE and Péclet number

Compression of the poroelastic gel induces a range of outward fluid fluxes under varying levels of stress and strain rates. This flux is characterized using the chemical potential gradient and permeability profiles obtained from the constitutive finite element modeling. FEM framework was used to simulate 30% compression of the gel over a 5-hour period, to represent the growth factor transport dynamics at a time scale comparable to the cellular growth time scale (∼ a few hours). Chemical potential as a function of radial position was recorded every 10 minutes during the first hour and then at hourly intervals for the remainder of the simulation (i.e., 2, 3, 4, and 5 hours). A logistic function was fitted to the chemical potential gradient data to characterize the radial profiles (eqn.14b). *L* represents the maximum attainable value of the chemical potential gradient in the domain (unit Pa), *k* is the steepness coefficient (unit 1/m) and *r*_0_ is the inflection point(unit m). Permeability was also extracted as a function of time from the FEM simulations, computed using (eqn.13).

The advection term *u*(*r, t*) associated with outward fluid flux was estimated using eqn.14a,14b.

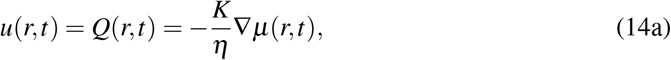

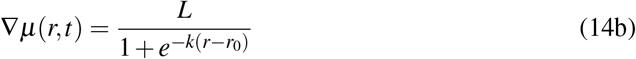

The fluid flux induced by compression influences the concentration profiles of diffusible substrates (e.g., nutrients, growth factors, metabolites) within the tumor microenvironment. To quantify this effect, we analyze growth factor concentration profiles within an advection-diffusion PDE framework, incorporating fluid flux as a governing transport mechanism in addition to diffusion.

The advection-diffusion equation was solved for an 8 mm *×* 8 mm two-dimensional square grid, designed to mimic the compression experiments on the alginate gel in silico. This grid represents a tumor microenvironment assumed to be isotropic and symmetric. In this PDE setup, we are not accounting for growth factor depletion due to consumption by tumor cells. We also do not account for heterogeneous perfusion, transvascular exchange or extravascular binding in the 2D tumor microenvironment. These terms can be accounted for using an additional reaction term in the PDE. Baxter and Jain provided an extensive theoretical framework to model advection-diffusion-reaction PDEs for macromolecules solute transport in the tumor microenvironment.^20,55–57^ This can be adapted for growth factors.

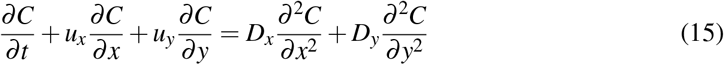

The advection-diffusion equation in two-dimensional Cartesian coordinates describes the transport of a scalar concentration field, *C*(*x, y, t*), under the combined effects of advection and diffusion (eqn.15). The spatial coordinates *x* and *y* represent length dimensions, while t denotes time. The velocity components in these directions are *u*_*x*_ and *u*_*y*_, respectively. The diffusion coefficients are assumed constant throughout the domain, with *D*_*x*_ = *D*_*y*_ = *D*.

To nondimensionalize the equation, characteristic scales were introduced based on the advection time scale: characteristic length *L*, velocity *U*, concentration *C*_0_, and advection time 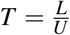. The resulting non-dimensional variables are defined as 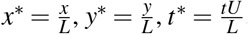, and 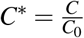, with velocity components 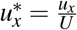 and 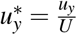.

The dimensionless Péclet number, the ratio between rates of advection to diffusion, given by *Pe* = *UL/D*, characterizes the relative influence of advection to diffusion on the characteristic length scale *L*. The convection-diffusion equation was non-dimensionalized using the advection time scale(eqn.16).

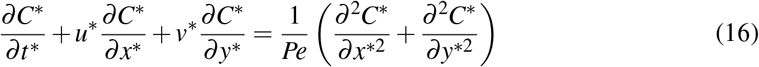

The characteristic length *L* is defined as 4 mm, consistent with the radial dimensions of the experimental setup used to study poroelastic effects under different conditions. The characteristic velocity *U* corresponds to the maximum radial velocity observed from the FEM simulations. The characteristic concentration *C*_0_ is set to a nominal growth factor concentration of 10^−7^M, reflecting typical concentration of growth factors in tumor microenvironment.

Diffusion coefficient of a growth factor (or other macromolecular solute in a tumor tissue or engineered gels) presented in literature typically ranges from 10^−8^ to 10^−12^m^2^*/*s, with variations influenced by the size, charge, porosity, and interaction with extracellular matrix components. (EGF in rat brain extra cellular space^58^: 10^−11^m^2^*/*s, VEGF in type 1 collagen hydrogel^38^: 10^−10^m^2^*/*s, IGF in fibrin gels^59^: 10^−8^m^2^*/*s). Interstitial fluid velocities in normal tissues are in the range 0.1 − 1 *µ*m/s, whereas velocities in tumor tissues have been noted to be as high as 55 *µ*m/s in some cases.^60^

#### 2. Numerical scheme for solving the PDE

The computational domain was defined over *x*^*∗*^, *y*^*∗*^ *∈* [0, 1] with symmetrical Dirichlet boundary conditions for growth factor concentration on all 4 edges of the grid. The initial condition was set as a uniform concentration of *C*^*∗*^ across the entire domain.

The advection-diffusion equation was discretized using the Method of Lines (MOL), with spatial derivatives approximated via finite-difference method on a uniform grid with spacing Δ*x*^*∗*^ = Δ*y*^*∗*^ = 0.05. The advection term was discretized using an upwind scheme (order 1) for numerical stability. The advection term is updated at discrete time intervals throughout the PDE numerical scheme to account for the change in chemical potential gradients and mobility with time. This update is done every 10 mins for the first hour and then, every hour, until the end of the simulation. Adjusting the frequency of updates to the advection term influences both the accuracy of the numerical solution and the computational efficiency. By updating more frequently during the initial phases of the simulation, we capture initially faster transient behaviors with greater precision.

Time integration was performed using the implicit second-order TRBDF2 (Trapezoidal Backward Differentiation Formula 2) method, which is suitable for stiff problems.

The numerical solution was implemented in Julia 1.11.3, utilizing ModelingToolkit.jl for symbolic modeling and equation generation, MOLFiniteDifference.jl for spatial discretization via the Method of Lines approach, and DifferentialEquations.jl for numerical solving of the advection-diffusion equation.

### H. Agent-based modeling for tumor growth under different growth factor availability

To investigate the impact of compression-induced growth factor gradients on tumor progression, an agent-based model (ABM) was developed using PhysiCell (Version 1.14.0)^61^, an opensource, physics-based cell simulator for multicellular systems. PhysiCell incorporates BioFVM^62^, a parallelized diffusive transport solver for 2D and 3D biological simulations, to simulate growth factor transport dynamics within the tumor microenvironment. Growth factor concentrations ranging from 0 to 10^−7^ M were used to parameterize the tumor microenvironment by specifying the initial and boundary conditions for substrate fields within the PhysiCell framework.

All agent-based simulations were executed on the Bridges-2^63^ cluster at the Pittsburgh Supercomputing Center. To capture stochastic variability, ten independent agent-based model (ABM) simulations were run per condition. Results are summarized as mean ± standard deviation. Between-condition differences in the means were evaluated with Welch’s two-sample t-test (*α* = 0.05, two-tailed), which does not assume equal variances. Snapshots of the simulation were saved as scalable vector graphics (SVGs) at specified time intervals for visualization.

The agent-based model was simulated on a 0.4 mm × 0.4 mm square domain (a small sub-section of the 8 *×* 8 mm alginate gel) with an initial tumor cell population of 200 cells. The simulation was run for 7 days (10080 minutes), with tumor cells proliferating in response to local growth factor concentrations. Each cell sampled its local microenvironment and determined its growth rate based on a sigmoidal relationship between growth factor concentration and cellular proliferation (Supplement S7).

Growth factor-receptor interactions typically exhibit saturable binding kinetics. This is depicted by a sigmoidal curve, where low concentrations result in minimal proliferation due to insufficient receptor activation, while increasing concentrations enhance proliferation rates up to a critical threshold. Beyond this point, receptor saturation limits further increase in proliferation despite higher extracellular growth factor levels.^64–66^ In our ABM, growth factor level cellular growth relationship was defined by a saturation growth rate of 10^−4^, a Hill coefficient of 2, and a half-max concentration of 10^−9^ M. These values were selected as an approximation for a generic growth factor but can be adjusted depending on the specific growth factor or available experimental data. Additional parameters used in the ABM were adapted from previously published PhysiCell simulations^61^ and are listed in the supplementary section (Table 1).

Previous computational studies have built agent-based and hybrid models to study the impact of vascular structure and function on emergent cell population behaviors^67^. These frameworks have also been used to understand how angiogenesis drives tumor progression and to model how vascular features influence drug delivery^68–70^.Since our ABM is designed as a computational platform to emulate cancer cell growth within an alginate gel geometry, we do not incorporate certain physiologically relevant factors, such as ECM heterogeneity, vascular heterogeneity, and nutrient competition, into the current model. Only heterogeneity in growth factor gradients generated by compression-driven interstitial flow is represented. Individual cancer cells are assumed to be phenotypically identical, and proliferation is governed solely by local growth factor concentration. Direct effects of solid stress, hydrostatic pressure, or matrix stiffness on cell cycle progression are not modeled. These simplifications help to isolate the effect of mechanical compression on growth factor transport and demonstrate its impact on tumor proliferation.

Future extensions of these models could couple multiscale mechanotransduction models, ECM remodeling, nutrient dynamics, and vascular heterogeneity to capture the joint influence of stress, stiffness, and biochemical cues on tumor growth.

## VI. SUPPLEMENTARY MATERIALS

Supplementary figures (S1-S9) and supplementary table 1.

## ACKNOWLEDGMENTS

This study was supported by the REgeneration and ReSToration of Functions in ORal HEalth (RESTORE) Prize funded by the Center for Innovation & Precision Dentistry (CiPD) and the Department of Oral and Maxillofacial Surgery’s Schoenleber Fund, at the University of Pennsylvania. KHV received support from the National Institutes of Health for this work, NIH/NIDCR R00DE030084. This work was partially supported by NSF through the University of Pennsylvania Materials Research Science and Engineering Center (MRSEC) (DMR-2309043) through a seed grant to KHV and PKP. PKP and YR acknowledge NSF grant DMR 2212162. PKP acknowledges partial support from NIH grant R01 HL148227. RR and SK acknowledge funding by the National Institutes of Health Grants R35GM136259, EB01775309 and U01CA250044. We acknowledge the use of the Bridges-2 system, a resource of the Pittsburgh Supercomputing Center (PSC), part of the ACCESS program under grant MCB200101.

## VII. ETHICS APPROVAL STATEMENT

Ethics approval is not required for these studies.

## VIII. AUTHOR CONTRIBUTIONS

Authors ZL, YR, SK, PKP, RR, PAJ, and KHV contributed to the conceptualization, data curation, writing, and review and editing of this manuscript. ZL, YR, SK, PM, and JK contributed to the investigation, data analysis, and visualization. All authors reviewed and approved final version of the manuscript.

## IX. CONFLICT OF INTEREST STATEMENT

The authors have no conflicts to disclose.

## X. DATA AVAILABILITY STATEMENT

The data that support the findings will be available in Dryad repository at [DOI to be provided] prior to publication.

## Notes

### Competing Interest Statement

The authors have declared no competing interest.

