## Supplemental Information for "Modeling tumor transport and growth with poroelastic biopolymer networks"

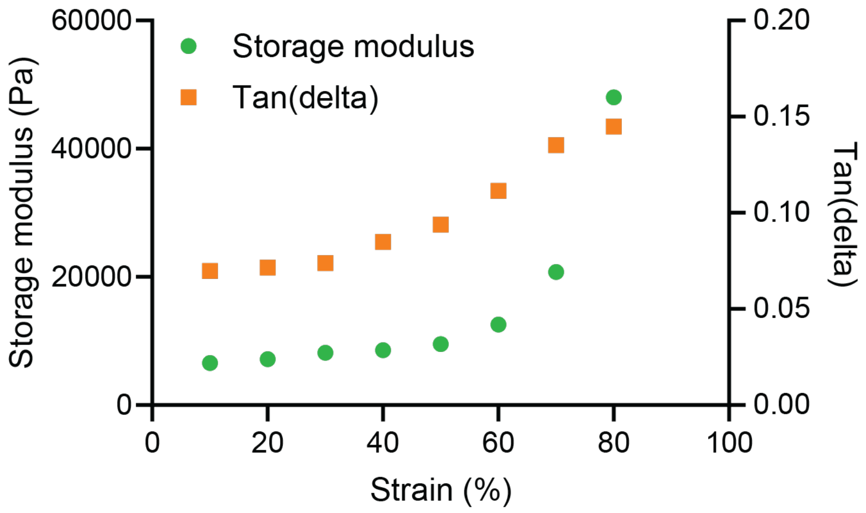


FIG. S1. Storage moduli and tan(delta) as a function of compressive strain for ionic Alg under slow stepwise compressions(strain rate 0.025%/s), where storage moduli and tan(delta) are measured when the stress is fully relaxed.

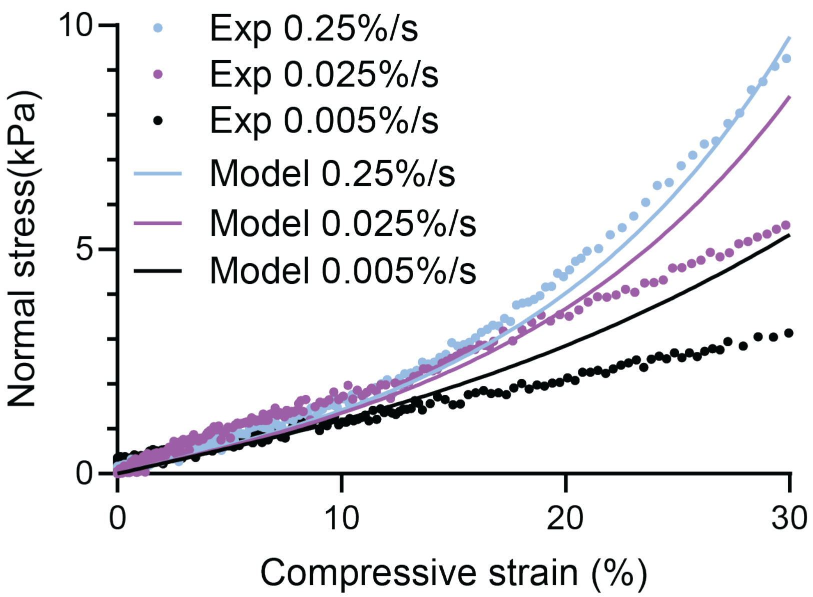


FIG. S2. Normal stress vs. compressive strain under various strain rates from experiments and FEM simulations.

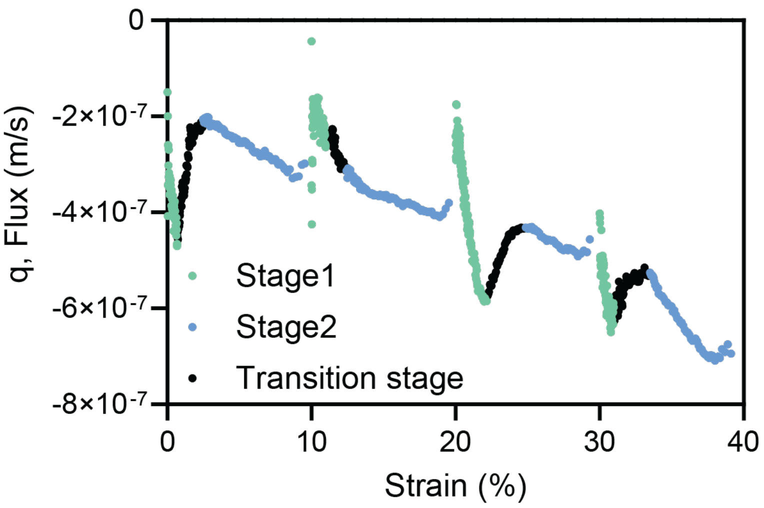


FIG. S3. Fluid flux vs. strain under stepwise compression, which repeatedly generated stage 1's fast-decreasing regions and the transition stage from stage 1 to stage 2.

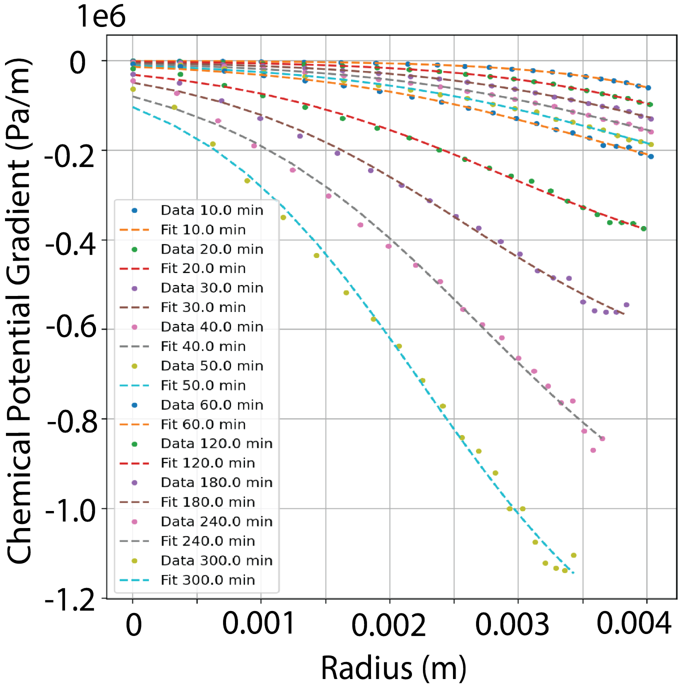


FIG. S4. 30 % compression in 5 hrs. The radial chemical potential gradient profiles were obtained from the FEM simulation results. This data was recorded at discrete times ($t$=10, 20,30,40,50,60,120,180,240,300 min) throughout the simulation. Logistic function was fit to radial chemical potential gradient profiles for all these time points. The gradients are relatively flat in radial direction at the initial time points. They become steeper as the simulation progresses.

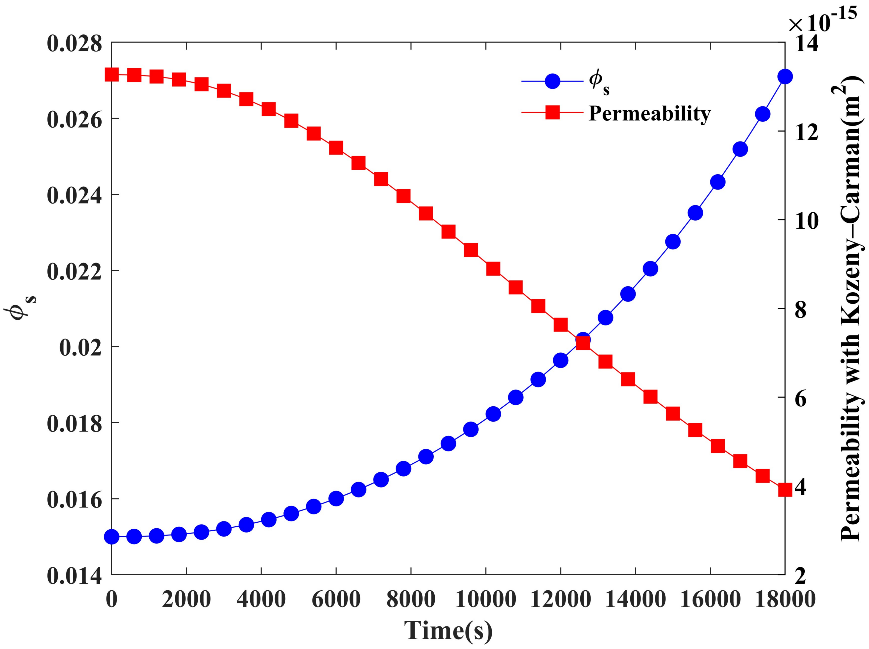


FIG. S5. Solid volume fraction and permeability values for the simulation lattice as a function of time, which is obtained by 30% compression in 5 hrs.

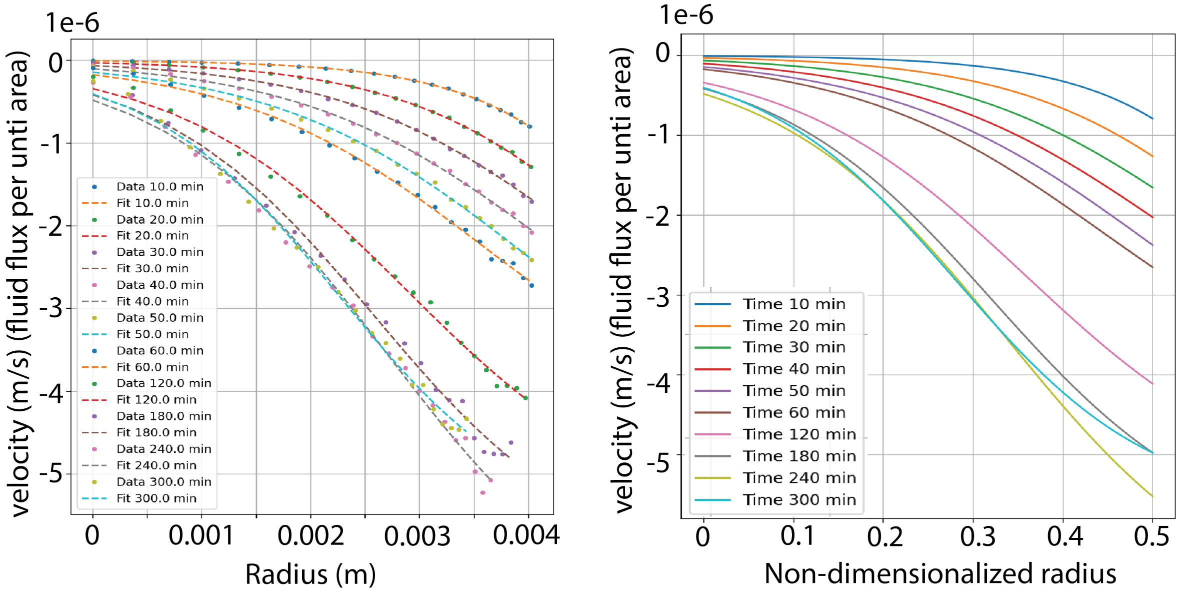


FIG. S6. Velocity is calculated as a product of chemical potential gradient and mobility. The first plot (left) shows the velocity profile (function of radius and time) as a function of time. The second plot (right) shows the mapped velocity profiles onto the non-dimensionalized radius for the advection-diffusion PDE setup.

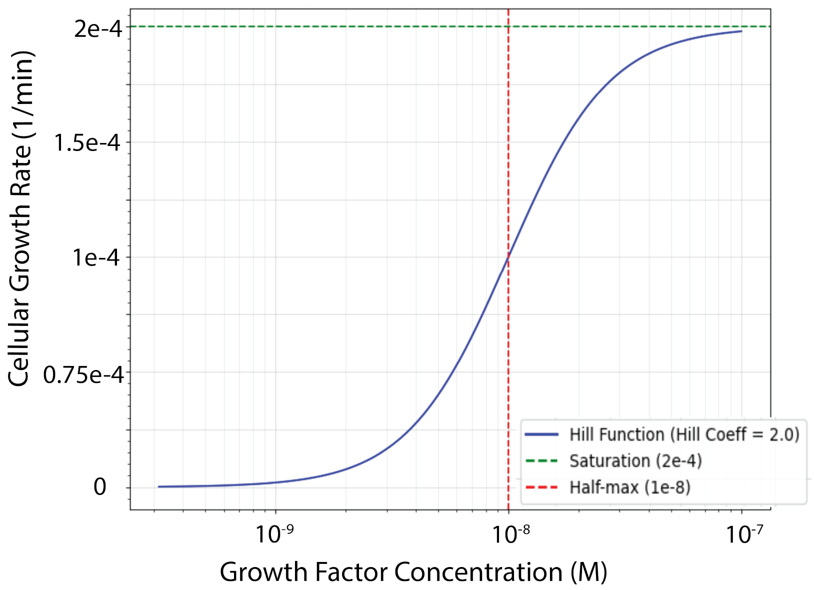


FIG. S7. The cellular proliferation rule in the ABM was set as a sigmoidal function of local growth factor concentration.

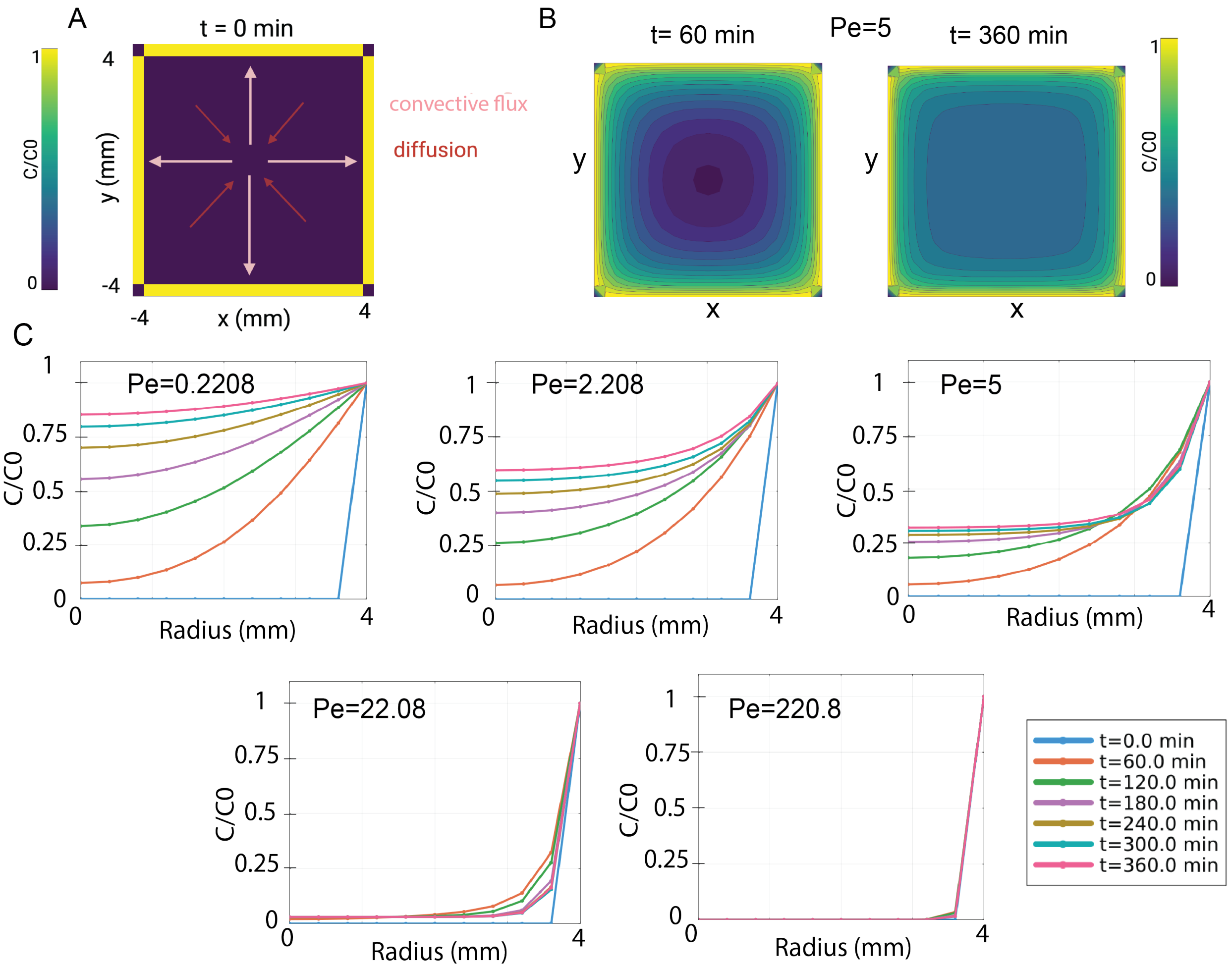


FIG. S8. Influence of Péclet number on growth-factor distribution - Simulations initialized from a Growth Factor Depleted Lattice. A) At $t$=0 min, the concentration field is 0 across the 8 mm×8 mm domain and boundary conditions are set to 1. Pink arrows indicate the imposed outward convective flux, while red arrows show the inward direction of diffusion. B.) Two-dimensional concentration contours at $t$=0 min (left) and $t$=360 (right) illustrate how the initially depleted field evolves under a moderate flow regime (here, Pe = 5.0). By 360 min, a nearly square-shaped plateau of low concentration persists in the interior, bounded by steep boundary layers. C.) Radial profiles of C/C0 at six time points (0, 60, 120, 180, 240, 360 min) for five Péclet numbers (0.2208, 2.208, 5.0, 22.08, 220.8). At low Pe, diffusion dominates and enables a significant influx of growth factors. As Pe increases, convective transport limits this inward flux and confines accumulation to narrow boundary regions.

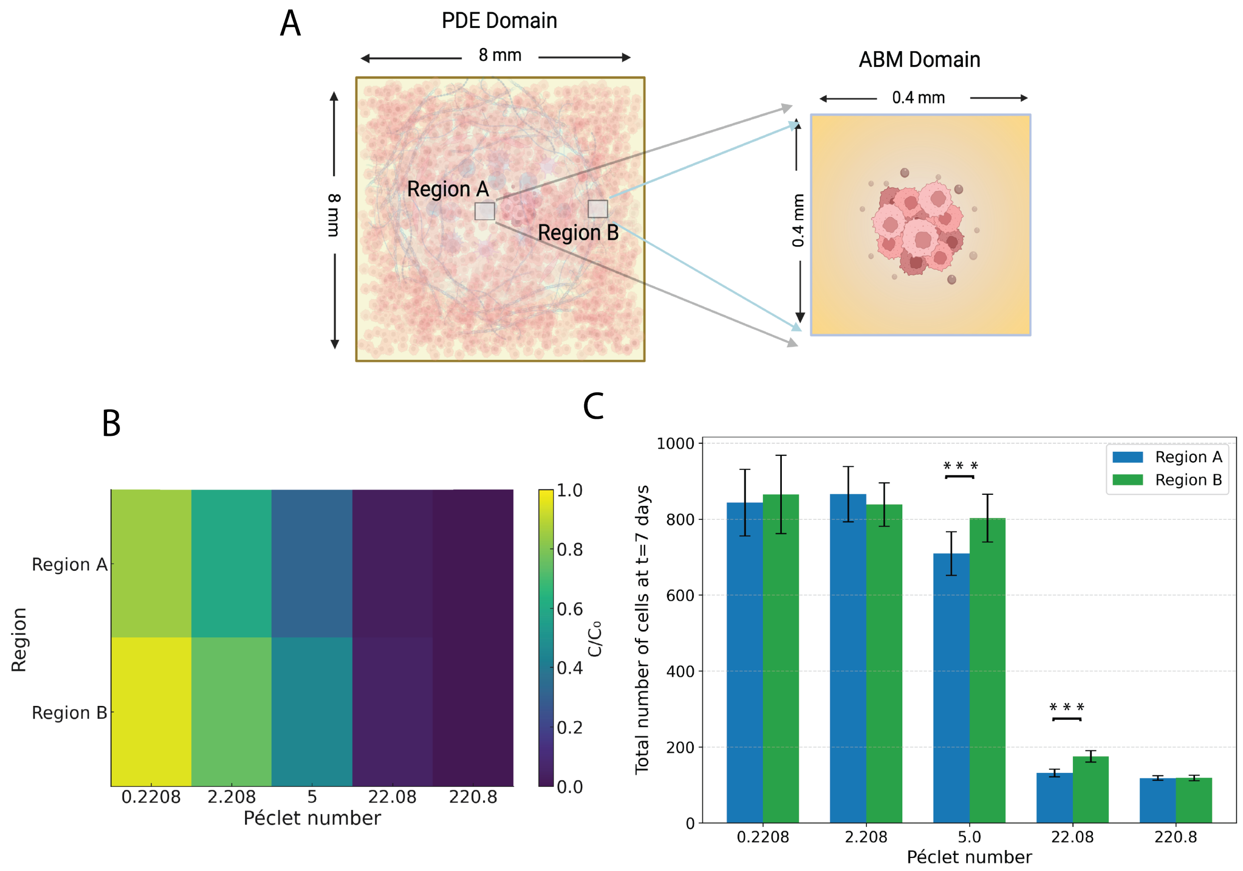


FIG. S9. Coupling growth factor transport and tumor growth results across scales A) Setup of the computational study - PDE domain on the 8 mm $\times$ 8 mm lattice and tumor growth ABM domain on 0.4 mm $\times$ 0.4 mm regions of the lattice. B) Heatmap - summarizing the Region A and Region B growth factor (GF) concentrations (at $t$=6 hrs). Spatial heterogeneity in growth factor availability between Regions A and B reduces as Pe increases. C) Tumor load at $t$ = 7 days / 10080 min) at Regions A and B, under different Pe. At intermediate Pe values, tumor growth differs significant between regions A and B. Tumor load bars show the $\mu \pm \sigma$ from ten independent ABM stochastic runs per condition. Welch's two-sample t-test was applied to each Region A-B pair; an asterisk indicates a significant difference. *(p<0.01)* and three asterisks indicate *p<0.0001*.

| **Parameter Description** | **Value** | **Unit** |
|---|---|---|
| Total simulation time | 7200 | min |
| Domain size in x-direction | 400 | µm |
| Domain size in y-direction | 400 | µm |
| Grid resolution (dx, dy) | 20 | µm |
| Initial number of cells (t=0) | 200 | cells |
| GF diffusion coefficient | 100 | $\mu m^2/s$ |
| GF decay rate | 0 | 1/min |
| GF initial condition | GF concentration from PDE solution | M |
| GF Dirichlet BC | GF concentration from PDE solution | M |
| Cell proliferation rate | $0–2 \times 10^{-4}$ (sigmoid function) | 1/min |
| Cell apoptosis rate | $5.3 \times 10^{-5}$ | 1/min |

TABLE I. Summary of ABM (PhysiCell) parameters used to generate a family of tumor growth curves from the agent based simulations.
